# Replacing *In Vivo* Experiments for PK/PD Target Determination Through *In Vitro* Time-Kill Experiments and PK/PD Modelling Incorporating Inter-strain Variability: Application to Meropenem Against *Pseudomonas aeruginosa*

**DOI:** 10.64898/2026.09.10.750598

**Authors:** Julien M. Buyck, Ombeline Krekounian, Tom Collet, Vincent Aranzana-Climent, Nicolas Grégoire

## Abstract

**Background:** Optimal antibiotic dosing regimens depend on the pharmacokinetic/pharmacodynamic (PK/PD) index that best predicts antibacterial efficacy. PK/PD targets are traditionally determined using murine infection models based on a limited number of bacterial isolates.

**Objective:** This study aimed to investigate whether animal experiments could be replaced by *in vitro* time-kill experiments performed on a large collection of clinical isolates and analyzed using a modelling approach accounting for inter-strain variability. The proposed framework was evaluated using meropenem against *Pseudomonas aeruginosa*.

**Materials and Methods:** *In vitro* time-kill experiments were performed on 66 clinical isolates of *P. aeruginosa*. A population pharmacodynamic model was developed from experimental data. A murine pharmacokinetic model was reproduced from literature and combined with the pharmacodynamic model to simulate *in vivo* bacterial burden over time. The relationships between simulated bacterial counts at 24 h and the three main PK/PD indices (fCmax/MIC, fAUC/MIC and %fT>MIC) were characterized using nonlinear mixed-effects Imax models.

**Results:** The PK/PD index showing the strongest correlation with meropenem efficacy at 24 h was %fT>MIC (R² = 0.989), compared with fAUC/MIC (R² = 0.373) and fCmax/MIC (R² = 0.284). These findings are consistent with previous studies using murine thigh infection models. The %fT>MIC target required to achieve a 2-log CFU reduction was estimated at 44%, with substantial inter-strain variability (10th and 90th percentiles: 27% and 71%, respectively).

**Conclusions:** Using meropenem against *P. aeruginosa* as a proof of concept, we demonstrate that *in vitro* time-kill experiments combined with pharmacometric modelling can identify the same PK/PD efficacy targets as animal infection models. Moreover, performing experiments on a large panel of clinical isolates enables the quantification of inter-strain variability in PK/PD targets, providing information that may improve their translation to clinical dosing optimization.

## Introduction

Antibiotics are commonly categorized based on the PK/PD index that best correlates with their antibacterial efficacy. They are thus described as concentration-dependent when efficacy is associated with the ratio of peak free drug concentration to the minimum inhibitory concentration (fCmax/MIC), as observed for aminoglycosides, or with the ratio of the free drug area under the concentration–time curve to the MIC (fAUC/MIC), as for fluoroquinolones. In contrast, time-dependent antibiotics exhibit efficacy linked to the fraction of the dosing interval during which free drug concentrations exceed the MIC (f%T>MIC), as is the case for β-lactams. Finally, some antibiotics display concentration-independent activity but with a sustained post-antibiotic effect, with efficacy also best described by fAUC/MIC, as for tetracyclines [1].

Depending on the antibiotic class, the most appropriate dosing regimens differ, for example continuous infusions for time-dependent antibiotics or intermittent administrations for concentration-dependent antibiotics. It is therefore essential to determine, for each antibiotic, the class to which it most closely relates and the PK/PD index target required to achieve efficacy. Preclinical determination of these indices and their targets can be performed either *in vitro* [2–4] or *in vivo* [2,5–7]. *In vivo* models consist of infection models, typically in neutropenic mice and require several dozen animals per bacterial strain [2,5–7]. In a context where efforts are made to reduce animal use in research [8], following the principles of replacement, reduction, and refinement (3Rs), it appears relevant to seek alternatives to *in vivo* experiments through *in vitro* approaches.

*In vitro* models rely on dynamic systems with time-varying antibiotic concentrations mimicking pharmacokinetic (PK) profiles. They are technically demanding, and preclinical PK/PD index determination is therefore typically conducted using only a limited number of strains (usually 4 to 5) from the target species [9]. This approach relies on the assumption that normalizing PK/PD indices by the MIC adequately captures potential inter-strain pharmacodynamic variability, such that a small panel of strains is sufficient to define both the relevant index and its associated targets. In contrast, *in vitro* time-kill experiments, performed at constant antibiotic concentrations, are simpler to implement than dynamic systems. It has been suggested that such data, when combined with pharmacodynamic modelling and simulation, can be used to infer PK/PD indices [10]. However, this strategy was established using experimental data from a single reference strain and does not account for possible variability between strains [11].

It can be hypothesized that determining PK/PD indices and their targets would be more relevant when accounting for inter-strain variability, using mixed effects pharmacodynamic models, and in particular by evaluating whether targets depend on MIC values or other specific characteristics (e.g., resistance genes). This approach is further facilitated by increasing the automation level of microbiological techniques for time-kill experiments, enabling their application across a large number of strains [12].

The evaluation of PK/PD indices for meropenem (MEM) against *Pseudomonas aeruginosa* (*P. aeruginosa*) has been performed using dynamic *in vitro* models [13], *in vivo* models [14], and PK/PD modelling based on time-kill data from two strains [15]. The index most strongly associated with MEM efficacy is f%T>MIC. In this study we assessed whether we could determine PK/PD efficacy targets for meropenem against *P. aeruginosa* by *in vitro* time kill experiments and PK/PD modelling incorporating inter-strain pharmacodynamic variability without performing *in vivo* experiments.

## Material and methods

The overall principle of the method is to simulate the expected outcomes of *in vivo* experiments in a neutropenic mouse thigh infection model using a mixed-effects pharmacodynamic model built from *in vitro* time-kill data. The *in vitro* mixed-effects PD model is built from time-kill experiments performed on a large number of bacterial strains, while the pharmacokinetic (PK) model in neutropenic mice is obtained from the literature. PK/PD indices and their associated targets are then determined by nonlinear mixed-effects modelling of simulated bacterial counts at 24 hours. This general framework is illustrated in **Figure 1**.

**Figure 1.**
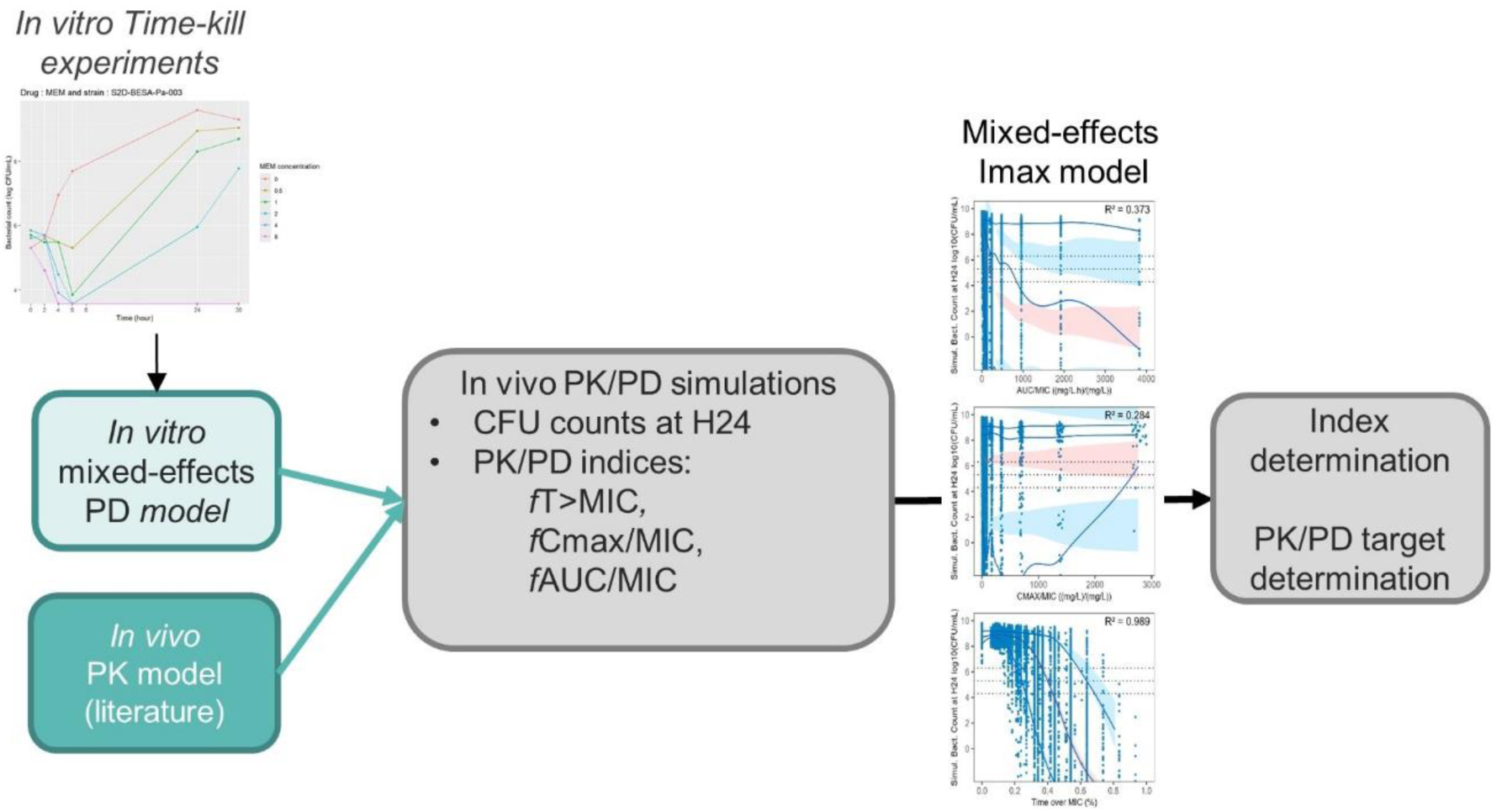
Flowchart of the method used to determine PK/PD indices from *in vitro* Time-Kill Curves.

### Meropenem in vitro degradation

The stability of MEM in MHB II broth at 35 ± 2 °C was investigated at five initial MEM concentrations: 0.5, 1, 2, 4 and 8 mg/L. MEM concentrations were measured at 7 time points over 96 h. Quantification of MEM concentrations was performed using a validated LC–MS/MS method as described previously [16].

### In vitro Time-Kill experiments

*In vitro* time-kill experiments (TKCs) were performed using a bacterial suspension from a 2 h logarithmic-growth-phase culture in Mueller–Hinton II broth (MHB II, Sigma-Aldrich, ref. 90922) was added to 24-well plate to achieve a final concentration of 1 x 10^6^ CFU/mL. A total of 66 *P. aeruginosa* clinical isolates were exposed to meropenem (MEM, Sigma-Aldrich, Saint-

Quentin-Fallavier, France, ref. PHR1772) at six predefined concentrations (0, 0.25, 0.5, 1, 2 and 4xMIC). The distribution of MIC values across clinical isolates is provided in **Figure S1**. Briefly, the plate was incubated at 35 ± 2°C for 30 h under agitation (130 rpm). Bacteria were sampled and Colony-forming unit (CFU) counts were measured at 0, 2, 4, 6, 24, and 30 h on MHA plate using spotting method [12] with a 48-Pin Microplate replicator (Boekel Scientific™, Feasterville, USA). The CFU number was then counted after incubation at 35°C for 16-20 h in an ambient air incubator. The limit of quantification (LOQ) was fixed at 1000 CFU/mL for 1 µL plated. One well without addition of antibiotic was included in each experiment as growth control. Two strains had MIC below the range of tested concentrations (≤0.5 mg/L) TKCs were designed as if they had an MIC of 0.5 mg/L for strain 457 and 0.25 mg/L for strain 494.

### In vitro mixed-effects pharmacodynamic model

The data were analyzed using the stochastic approximation expectation–maximization (SAEM) algorithm implemented in Monolix [17].

MEM degradation data were modelled independently from time-kill data. MEM degradation was described using a mono-exponential decay model with a first-order rate constant (Kdeg). The estimated Kdeg value was subsequently fixed during the modelling of the time-kill data.

The model was fitted to log10-transformed CFU data. Observations were assumed to follow a normal distribution with a constant error model. Measurements below the limit of quantification (LOQ) were handled as censored data using Beal’s M3 method [18].

The structural model for the time-kill data included three components: a bacterial growth model, a MEM effect model, and an adaptation model to account for bacterial regrowth observed in some conditions.

For the growth component, the typical initial inoculum was fixed to the theoretical value (Inoc=10⁶ CFU/mL), adjusted by an estimated multiplicative factor (p). Several growth models were evaluated, including logistic growth (with a maximum bacterial load, Bmax), with or without a lag time, as well as a model incorporating an initial dormant bacterial state (B_R_) transitioning to a proliferating state (B_S_) through a first-order rate constant (Krs).

The drug effect was characterized using linear, power, Emax, and sigmoidal Emax models. Adaptive resistance was modeled using linear, power, Emax, and sigmoidal Emax functions applied to the parameters of the MEM effect model.

Inter-strain variability was modeled using log-normal distributions (except normal distribution on log10(B_max_), log10(Inoc), and the γ exponents of power functions, and logit-normal distribution for f, the corrective factor for MEM effect on the dormant bacterial subpopulation B_R_). Variability terms estimated to be negligible (coefficient of variation < 5%) were fixed to zero. Similarly, variability terms associated with relative standard errors (RSE) greater than 50% were removed. Potential correlations between random effects were also investigated.

The influence of MIC on selected structural parameters (excluding exponent terms) was evaluated. When MIC values were below the tested concentration range, they were set to half the lowest tested concentration. The relationships tested between MIC and model parameters included log-linear, linear, and power functions.

Model selection was based on the Akaike Information Criterion (AIC), and the final model was evaluated using standard diagnostic plots [19]. A nonparametric bootstrap was performed using Monolix default settings with 1000 replicates of datasets comprising the 66 clinical isolates. Model robustness was assessed by comparing parameter estimates from the final model with the median values obtained from the bootstrap analysis. Parameter precision was further evaluated by calculating relative standard errors (RSEs) from the bootstrap estimates, in addition to those derived from the stochastic approximation implemented in Monolix

### In vivo PK/PD dose fractionation studies simulations

A PK model described by Katsube *et al.* was used to simulate unbound plasma concentration profiles of MEM in mice [13]. This model was a one-compartment model with a depot compartment, an absorption rate constant K_a_ = 10.2 h⁻¹, a clearance CL = 0.033 L/h, a volume of distribution V_d_= 0.014 L, and a free fraction fu = 0.81.

*In vivo* bacterial count outcomes were simulated by combining the previously described *in vitro* PD models and *in vivo* PK models with Simulx [20]. The initial bacterial concentration (CFUs) used for the simulations was that reported in the original article at the time of MEM treatment initiation (log10_Inoc_ = 6.3 CFU/thigh and p = 1) [14]. The estimated maximum bacterial load value obtained *in vitro* was adjusted to the *in vivo* data from the control group of the original study (log10_Bmax_ = 8.76). Furthermore, in the *in vivo* PD model, at the time of MEM administration (2 hours after *P. aeruginosa* inoculation), all bacteria were assumed to be in the growth phase. The simulations were performed without accounting for residual variability.

CFUs were simulated at 24 hours for 12 dosing regimens reproducing those used in the study by Sugihara *et al.* [14], namely: a control group, 50 mg q3h, 100 mg q6h, 200 mg q12h, 400 mg q24h, 100 mg q3h, 200 mg q6h, 400 mg q12h, 800 mg q24h, 200 mg q3h, 400 mg q6h, and 800 mg q12h. The *in vivo* outcomes were simulated for 990 different bacterial strains (15 simulations for each of the 66 clinical isolates), with each isolate subjected to the 12 different dosing regimens (resulting in a total of 11,880 simulated profiles).

### Mixed-effects Imax model for PK/PD indices determination

The simulated data were modeled using a mixed-effects model with the SAEM algorithm with Monolix [17]. The simulated CFU counts at 24 hours were modeled as a function of f%T>MIC, fCmax/MIC, and fAUC/MIC using the following sigmoidal Imax model:

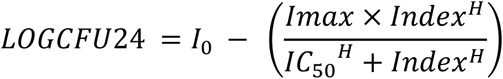

where LOGCFU24 is the bacterial load at T = 24 h; I₀ is the LOGCFU24 in the absence of treatment; Imax is the maximum reduction in bacterial load at 24 hours; Index is the value of the PK/PD index of interest (f%T>MIC, fCmax/MIC, or fAUC/MIC); H is the Hill coefficient; and IC₅₀ is the index value at which the reduction in bacterial load equals Imax/2. The effect of MIC on the parameters of this model was tested based on AIC values. Correlations between model parameters were estimated.

The PK/PD index that best predicted MEM efficacy was determined as the one with the highest coefficient of determination (R²) from this regression. Target values for this index were established for bacterial load reductions of 0 (bacteriostasis), 1, and 2 log₁₀(CFU/thigh).

### Bibliographic search for in vivo MEM PK/PD studies

A bibliographic search was performed to find *in vivo* data to validate our predictions. Four PubMed requests were used (**Text S1**) to find publications with PK/PD studies of MEM against *P. aeruginosa* strains with an MIC ≤ 32 µg/mL in a neutropenic mice thigh infection model. Publications were selected if they reported PK/PD targets and/or CFU/thigh (or ΔCFU/thigh) observed after MEM exposure linked with PK/PD index values. Data were extracted directly from tables and text or digitized from figures using WebPlotDigitizer v5.2 [21]

## Results

### Meropenem in vitro degradation half-life was 35h

The first-order degradation rate constant of MEM was estimated at 0.02 h⁻¹ (degradation half-life of 35 h) (**Figure 2**).

**Figure 2.**
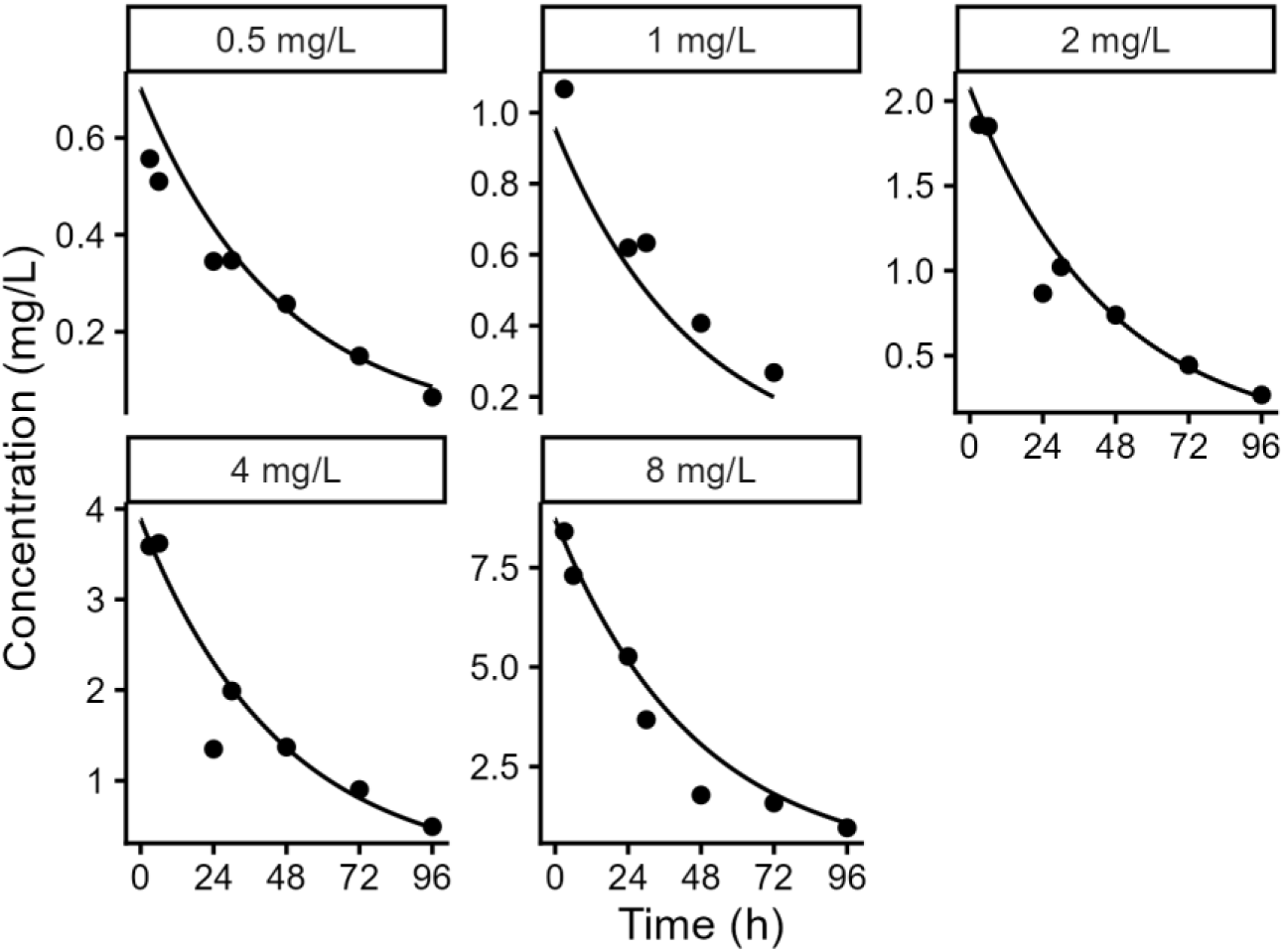
Meropenem *in vitro* degradation data and model fit. Points correspond to measured meropenem concentrations; lines correspond to mono-exponential model fit. One panel per initial concentration.

### In vitro time-kill data was adequately described by mixed-effects pharmacodynamic model

The *in vitro* pharmacodynamic model structure is presented in **Figure 3**.

**Figure 3.**
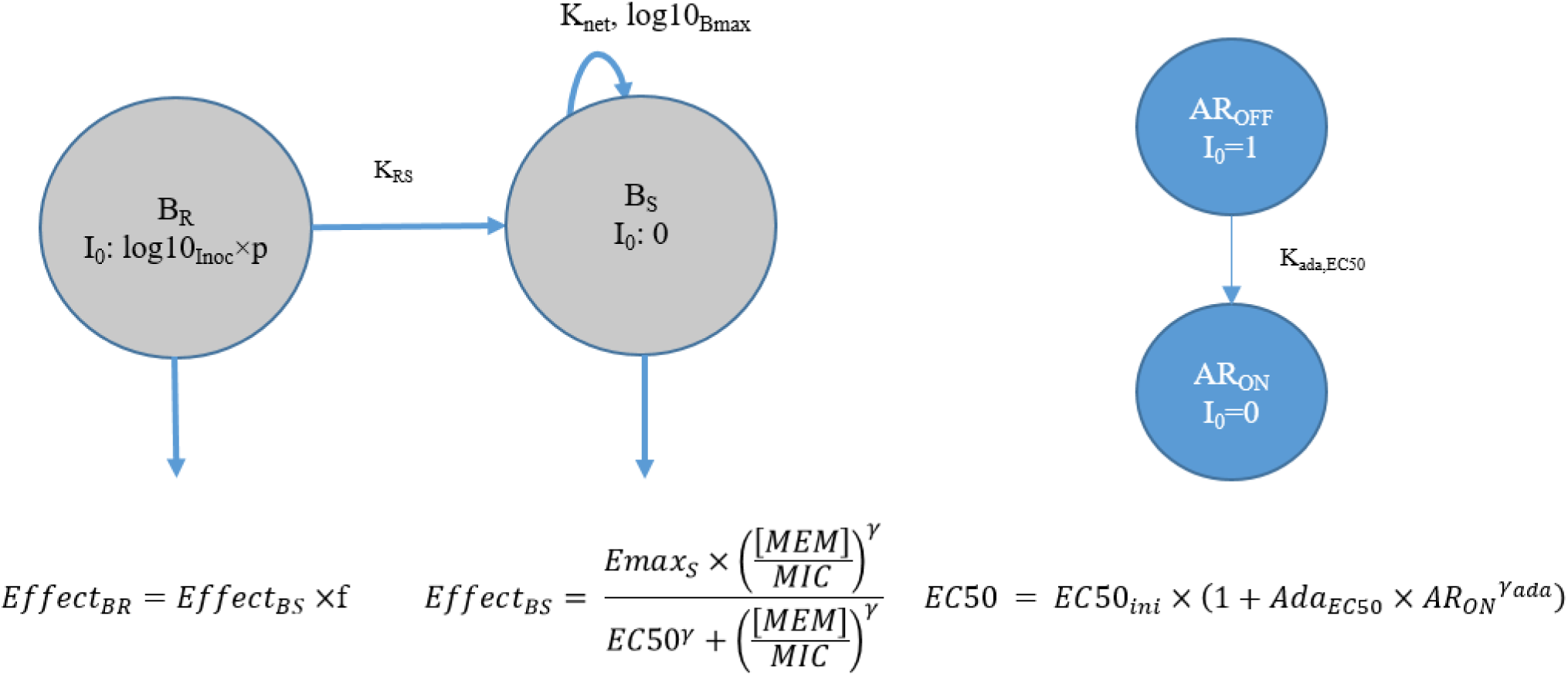
Graphical representation of the *in vitro* pharmacodynamic model. The growth and MEM effect model is shown in grey, and the adaptation model in blue.

Bacteria were initially in a non-replicative state (B_R_), with the initial inoculum deviating from the theoretical inoculum by a factor p = 0.5. Transition to a replicative state (B_S_) occurred with a first-order rate constant (K_RS_), and bacterial replication in the B_S_ state followed first-order kinetics (K_net_). A maximum bacterial burden (log10_Bmax_) was estimated. The effect of MEM was described by a sigmoidal model with parameters Emax_S_, EC50, and γ (see Equation 1 in Table 1). The effect of MEM on dormant bacteria compared with replicating bacteria was characterized by a factor f. Bacterial susceptibility decreased over time according to an adaptation model. Transition from the non-adapted state (AR_OFF_) to the adapted state (AR_ON_) followed a first-order rate constant (K_ada,EC50_). EC50 increased as a function of the AR_ON_ state according to a power model with parameters EC50_ini_, Ada_EC50_, and γ_ada_ (see Equation 2 in **Table 1**).

**Table 1.**
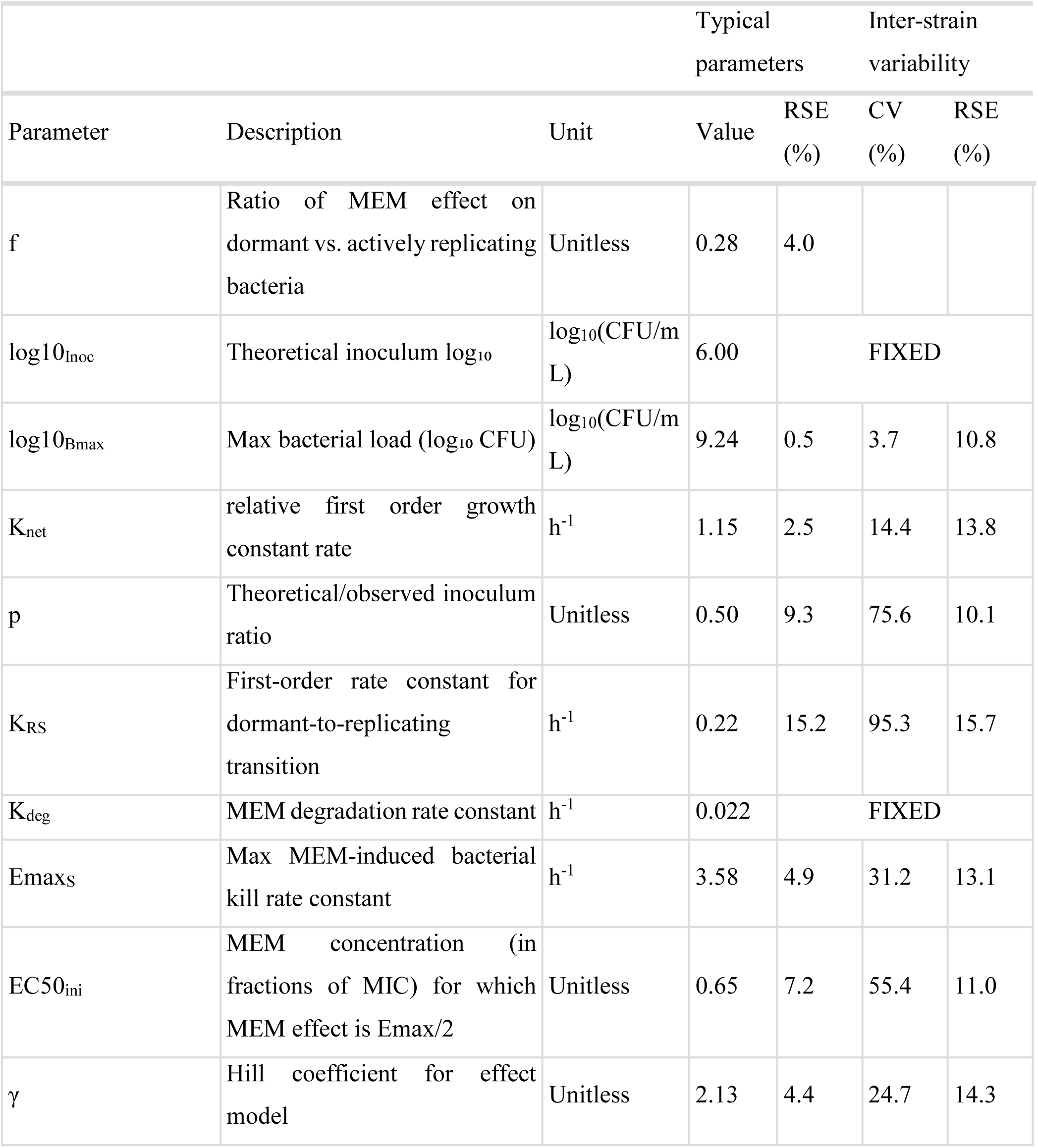
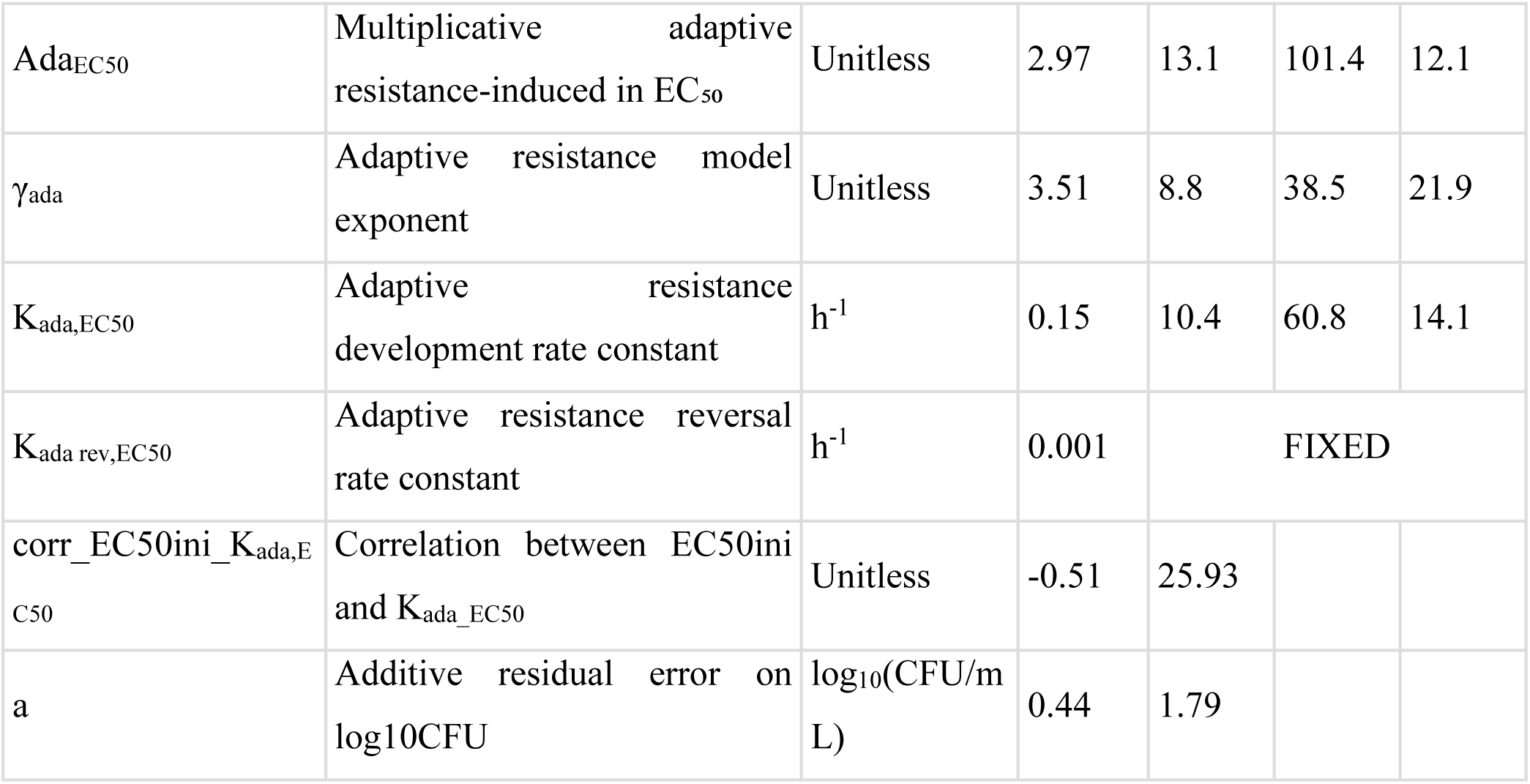
Final *in vitro* pharmacodynamic model parameters.

Inter-individual (inter-strain) variability was estimated for log10_Bmax_, K_net_, p, K_RS_, Emax, EC50_ini_, γ, Ada_EC50_, and γ_ada_. A significant negative correlation was observed between EC50_ini_ and K_ada,EC50_, indicating that lower EC50_ini_ values were associated with faster adaptation. Parameter estimates for the final model are presented in **Table 1**.

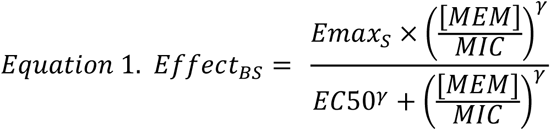

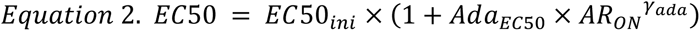

A representative example of time-kill experiment results for a single strain, along with predictions from the final *in vitro* pharmacodynamic model, is shown in **Figure 4**. Individual plots for all strains are presented in **Figure S2**.

**Figure 4.**
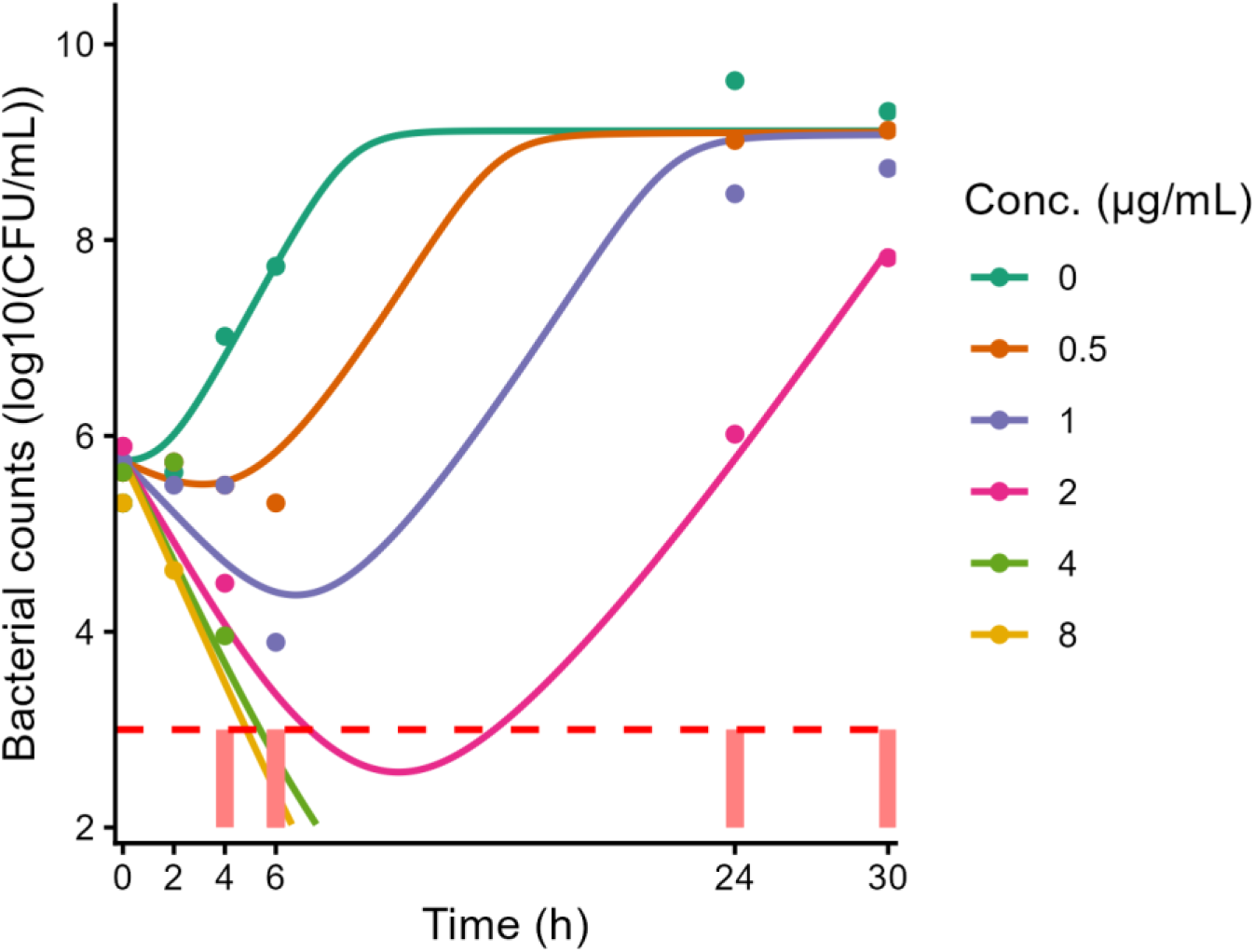
Representative time-kill data (points) measured for strain 003 (MIC = 2 mg/L), with final model predictions (lines). Red dashed horizontal line represents the limit of quantification (1000 CFU/mL). Light red rectangles signify that some data was below the limit of quantification.

Visual predictive checks are presented in **Figure 5**. Normalized prediction distribution errors and plots of observations versus typical and individual predictions are shown in **Figures S2– S5**. Overall, goodness-of-fit diagnostics did not indicate model bias, and parameter estimates from the final model were robust to resampling (bootstrap, **Table S1**).

**Figure 5.**
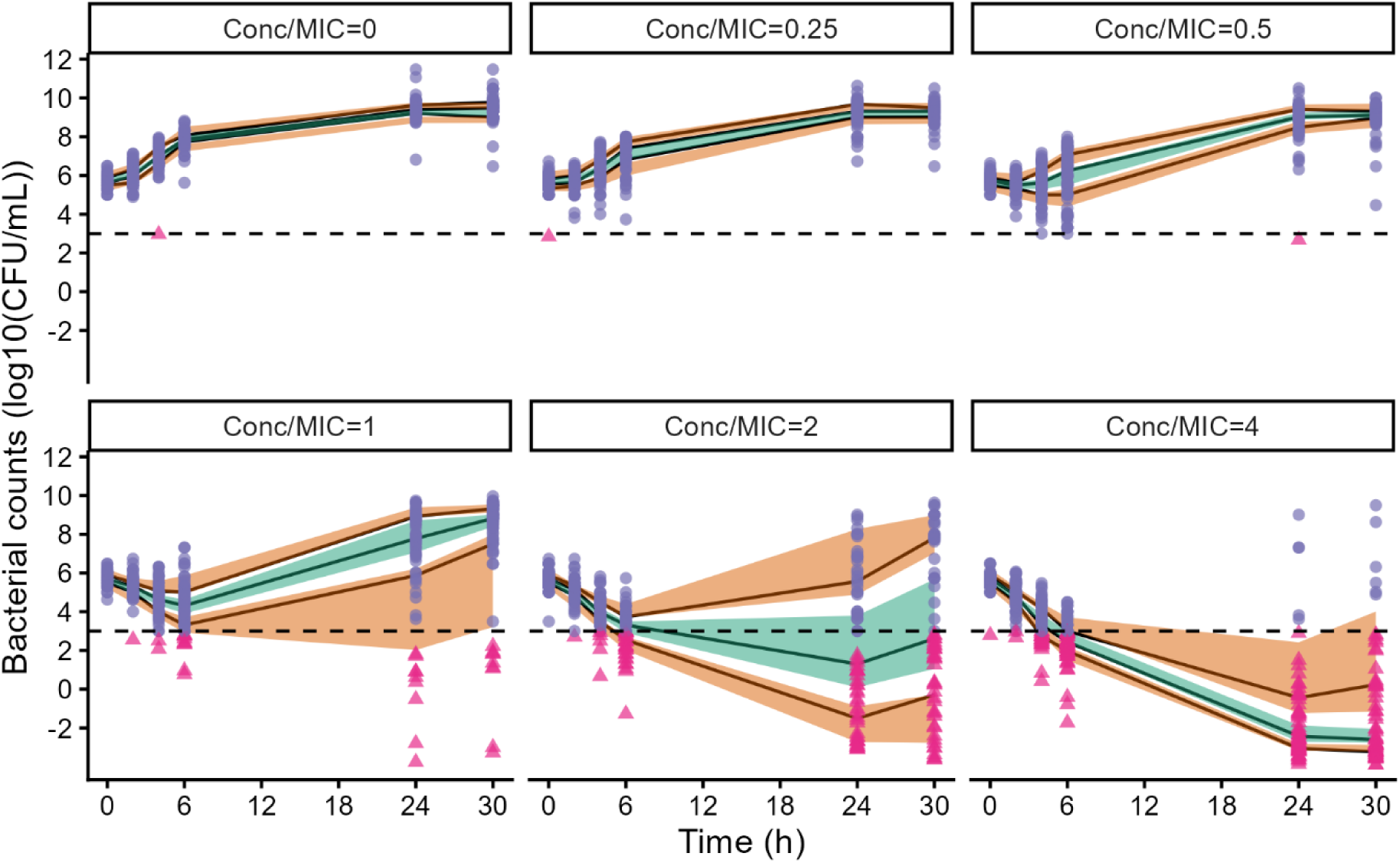
Visual predictive checks (VPCs) stratified by concentration-to-MIC ratios. Points represent observations (orange triangles indicate predictions for observations below the limit of quantification), lines denote the 25th, 50th, and 75th percentiles of the observations, and shaded areas show the 90% confidence intervals of the predicted 25th, 50th, and 75th percentiles (orange for the 25th and 75th percentiles, green for the 50th percentile). The dashed horizontal line represents the limit of quantification: 1000 CFU/mL.

In **Figure 5**, low variability in CFU counts over time can be observed for the control (concentration/MIC = 0), whereas substantially greater variability is seen when the concentration-to-MIC ratio equals 1. To illustrate this inter-strain variability, at MEM concentrations equal to the MIC, at 30 h, 9 measurements were below the limit of quantification (10³ CFU/mL) and 53 exceeded 10⁶ CFU/mL. At 4× MIC, at 30 h, measurements were below the limit of quantification in 58 cases and above 10⁶ CFU/mL in 3 cases.

### f%T>MIC was correctly identified as the PK/PD index most correlated with efficacy without performing in vivo experiments

Results from a simulated *in vivo* experiment for a *P. aeruginosa* strain following subcutaneous MEM dosing at 50, 100, 200, 400, and 800 mg/kg administered q24h, q12h, q6h, and q3h (selected from [14]) are shown in **Figure 6**. Median results across different MIC values are presented in **Figure S6**.

**Figure 6.**
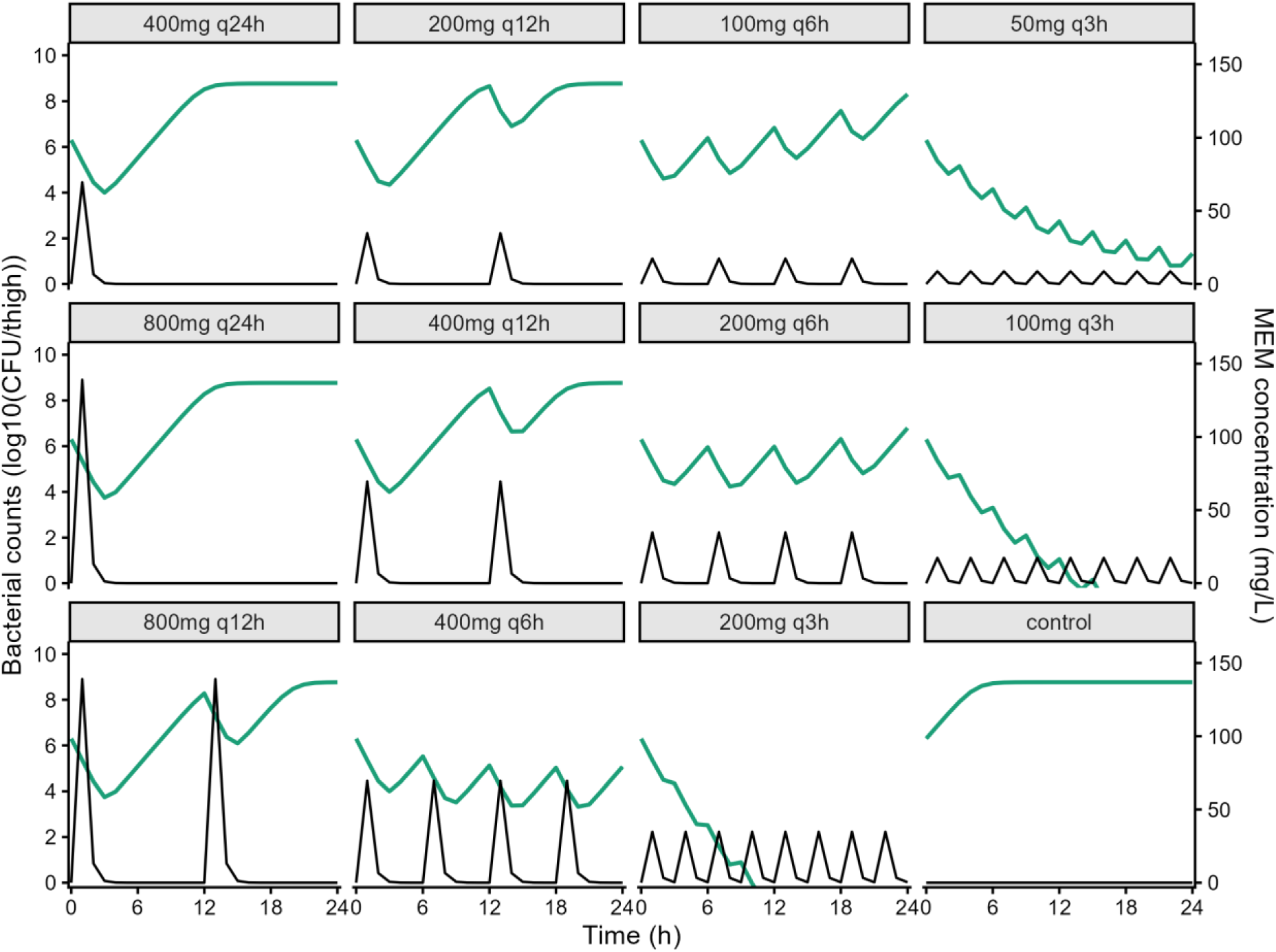
*In vivo* simulations of bacterial counts and plasma MEM concentrations following subcutaneous MEM dosing for a simulated isolate (Simulated strain #1, MIC = 2 mg/L) at 0 (control), 50, 100, 200, 400, and 800 mg/kg q24h, q12h, q6h, and q3h. Bacterial counts are shown in green (left axis) and plasma concentrations in black (right axis). First row: 400 mg total daily dose, Second row 800 mg total daily dose, Last row 1600 mg total daily dose + control.

Individual predictions from nonlinear mixed-effects Imax models for PK/PD index calculations for a representative strain (Simulated strain #1, MIC = 2 mg/L) are shown in **Figure 7**. Overall results of these PK/PD index analyses across all simulations are presented as VPCs in **Figure 8**.

**Figure 7.**
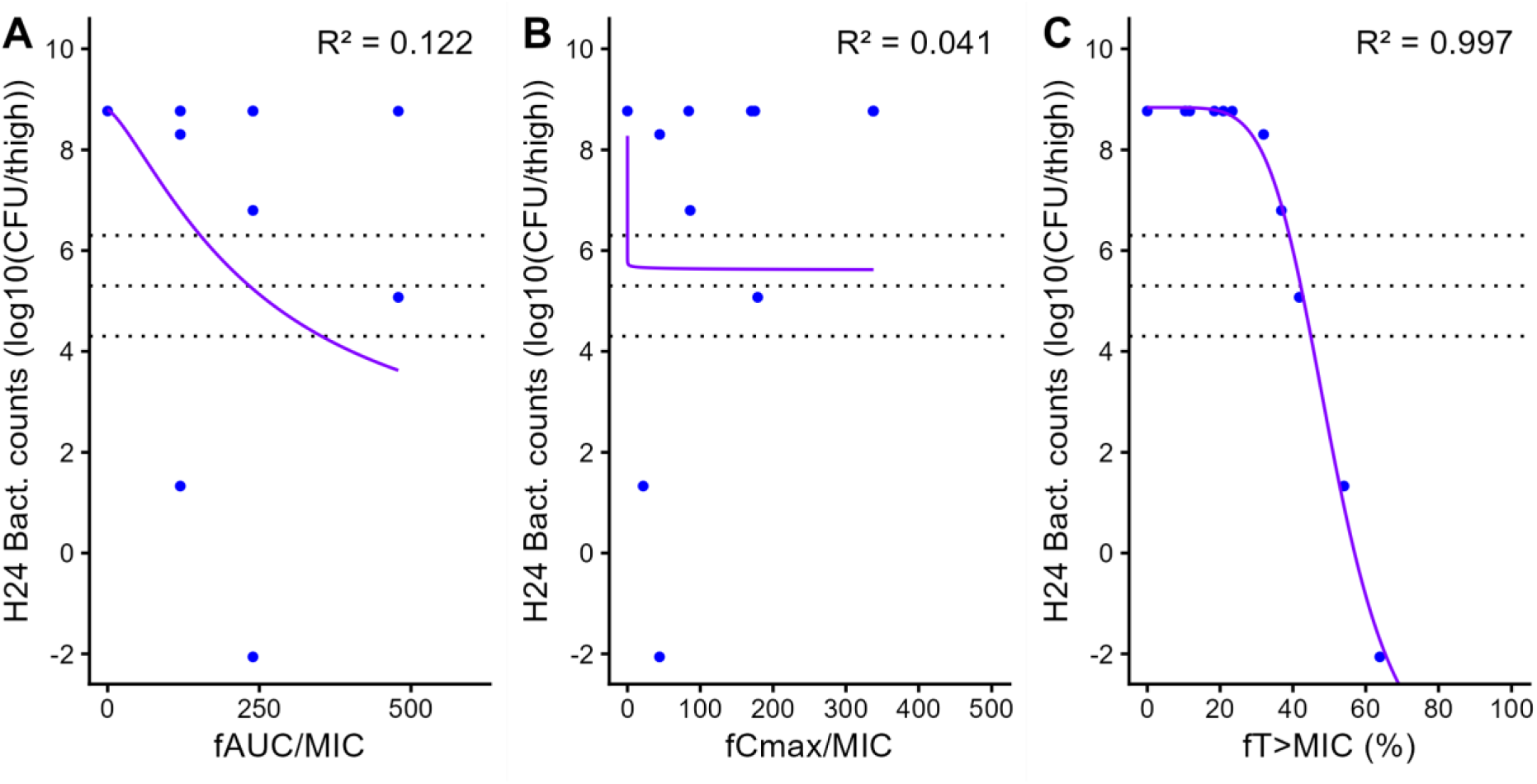
Individual predictions of bacterial counts at 24 h from Imax models for different PK/PD indices (A-fAUC/MIC;B-fCmax/MIC;C-f%T>MIC) for a simulated isolate (Simulated strain #1, MIC = 2 mg/L). Points represent bacterial burdens simulated by the *in vivo* PK/PD model for PK/PD index values corresponding to the 12 dosing regimens. Dotted lines represent stasis, 1logkill and 2logkill (from top to bottom).

**Figure 8.**
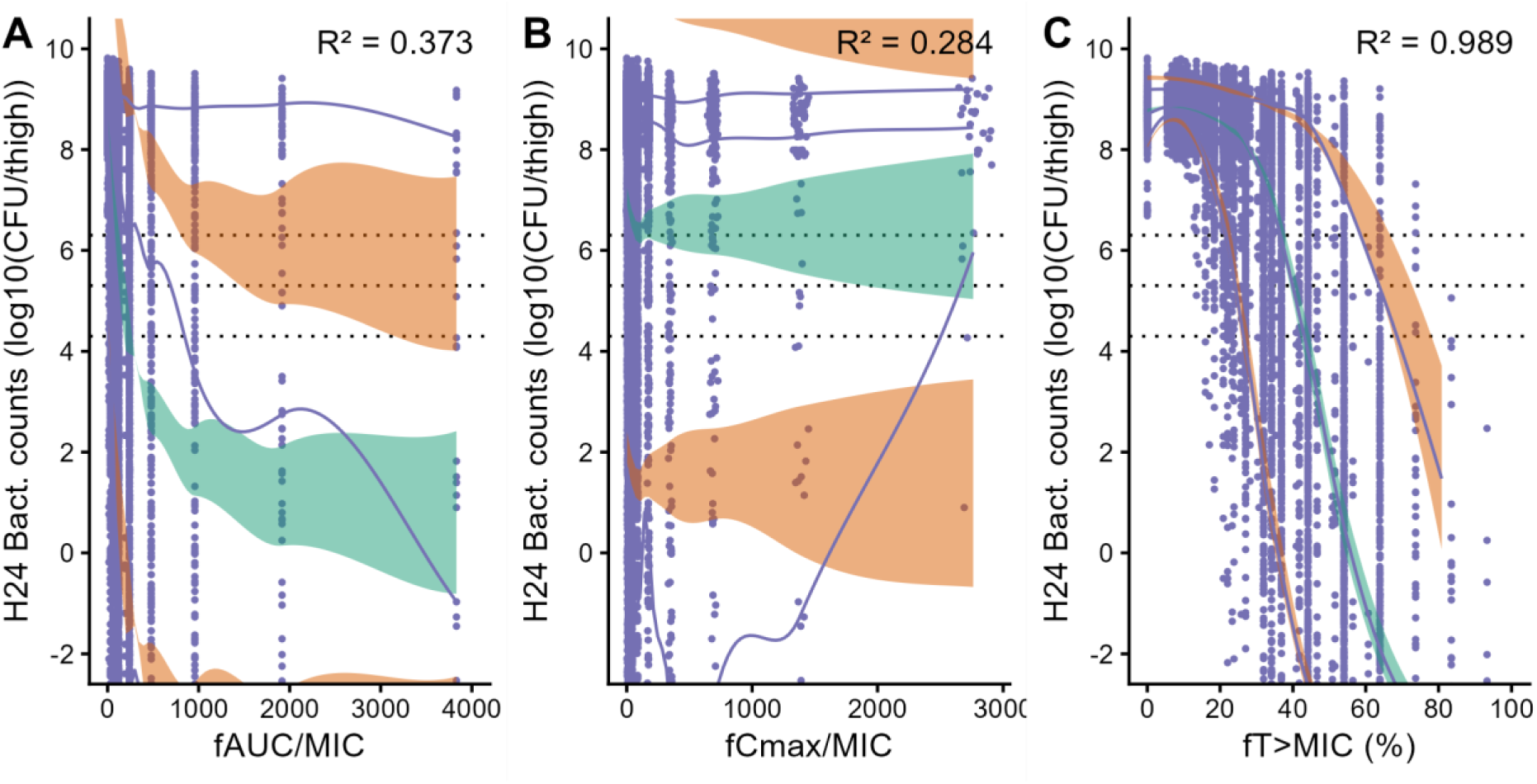
Visual predictive check of Imax models for different PK/PD indices (A-fAUC/MIC;B-fCmax/MIC;C-f%T>MIC) across the 990 simulated strains. Points represent simulated bacterial counts; LOESS curves denote the 10th, 50th, and 90th percentiles of the simulated data; shaded areas show the 90% confidence intervals of the predicted 10th, 50th, and 90th percentiles from the Imax model (orange for the 10th and 90th percentiles, green for the 50th percentile).

Coefficients of determination (R²) for fAUC/MIC, fCmax/MIC, and f%T>MIC were 0.373, 0.284, and 0.989, respectively. The 10th, 50th, and 90th percentiles of target f%T>MIC values required to achieve 0, 1, and 2 log10 CFU reductions at 24 h were (0.23, 0.38, 0.63), (0.25, 0.41, 0.68), and (0.27, 0.44, 0.71), respectively.

No significant covariate effect of MIC on different model parameters was found, showing that the PK/PD relationship was not different between low MIC and high MIC strains after normalization by MIC. Parameter estimates for the Imax model based on f%T>MIC are presented in **Table 2**.

**Table 2.** Imax mixed-effects model parameters for the f%T>MIC index.

| Parameter | Description | Unit | Typical parameter |  | Inter-strain variability |  |
| --- | --- | --- | --- | --- | --- | --- |
|  |  |  | Value | RSE (%) | CV (%) | RSE (%) |
| I<sub>0</sub> | LOGCFU24* without treatment | log<sub>10</sub>(CFU/thigh) | 8.8 | 0.1 | 3.6 | 2.6 |
| I<sub>max</sub> | Maximum decrease of LOGCFU24 | log<sub>10</sub>(CFU/thigh) | 13.5 | 0.4 | 6.3 | 5.4 |
| IC<sub>50</sub> | f%T>MIC value for which LOGCFU24=I<sub>0</sub>-I<sub>max</sub>/2 | f%T>MIC | 0.49 | 1.1 | 37.1 | 2.6 |
| H | Hill coefficient | Unitless | 5.69 | 0.9 | 22.5 | 3.5 |
| Corr I<sub>0</sub>_H | Correlation between I<sub>0</sub> and H | Unitless | -0.32 | 11.8 |  |  |
| Corr IC<sub>50</sub>_H | Correlation between IC<sub>50</sub> and H |  | 0.55 | 5.5 |  |  |
| Corr I<sub>max</sub>_H | Correlation between I<sub>max</sub> and H |  | -0.64 | 7.9 |  |  |
| Corr IC<sub>50</sub>_I<sub>0</sub> | Correlation between IC<sub>50</sub> and I<sub>0</sub> |  | -0.11 | 33 |  |  |
| Corr I<sub>max</sub>_I<sub>0</sub> | Correlation between I<sub>max</sub> and I<sub>0</sub> |  | 0.47 | 10.1 |  |  |
| Corr I<sub>max</sub>_IC<sub>50</sub> | Correlation between I<sub>max</sub> and IC<sub>50</sub> |  | -0.11 | 60.2 |  |  |
| a | Additive residual error on LOGCFU24 | log<sub>10</sub>(CFU/thigh) | 0.36 | 0.8 |  |  |

\* LOGCFU24 is the log<sub>10</sub> bacterial count at T=H24

### Predicted PK/PD efficacy targets were compatible but slightly higher than literature data

Our PubMed search was last updated on 2025-06-24 and unveiled 3 articles [22–24] with PK/PD targets (7 strains total). Data extracted from these articles is available in supplemental material.

**Figure 9** shows f%T>MIC targets to reach stasis, 1logkill and 2logkill according to our simulations compared with values extracted from the literature. Interestingly, the simulated distributions of %fT>MIC are right-skewed, almost log-normal. Our simulations might underestimate the effect of MEM for stasis and 1logkill, since all literature targets are between the 2.5^th^ and 50^th^ percentiles of the simulated distribution. For 2logkill, literature targets are centered around the median of the simulated distribution.

**Figure 9.**
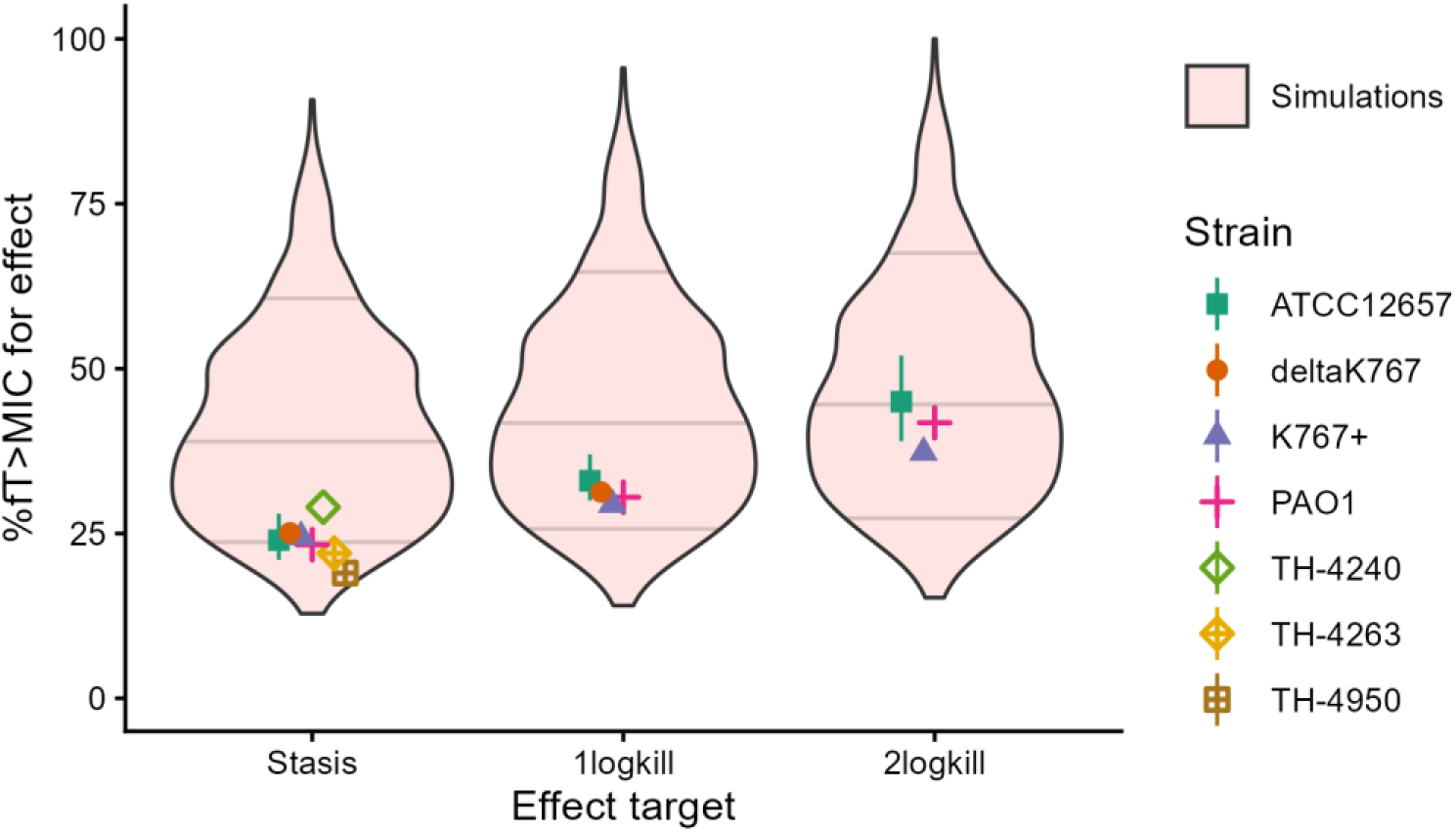
Violin plot comparing model simulated %fT>MIC targets to literature values. Light pink area represents model based 95% prediction interval of %fT>MIC targets necessary to reach stasis, 1logkill and 2log kill. Horizontal lines represent the 10^th^, 50^th^ and 90^th^ percentiles of the simulations. Dots represent literature medians and error bars represent reported 2.5^th^ and 97.5^th^ percentiles when reported.

However, the literature targets stay within the 95^th^ prediction interval of our simulations and given the low number of published data, the apparent underestimation of MEM effect could be random.

## Discussion

Using the approach applied in this study—combining PK/PD mixed-effects modeling of *in vitro* bactericidal data, simulation of *in vivo* mouse experiments, and subsequent PK/PD index determination—we identified f%T>MIC as the index most strongly associated with MEM efficacy. Median target values required to achieve a static effect, and 1- and 2-log10 CFU reductions were 38%, 41%, and 44%, respectively. This is compatible with data from [22–24] as shown on **Figure 9**. Interestingly, literature targets for stasis are in the bottom 25% of our prediction interval while literature targets for 2logkill are close to the median of our prediction interval. Our model thus doesn’t predict precisely what happened in *in vivo* thigh infection experiments. This could be due to multiple causes including: systematic differences between the literature strains and our strains; low sample size or even differences in bacterial environment making the bacteria respond differently to MEM. One could attempt to engineer TKC medium able to reproduce more closely the *in vivo* environment (as opposed to rich MHB-II medium) to see whether bacteria respond differently to MEM treatment and develop a model able to describe this data. However, despite these differences, our method was able to predict the correct PK/PD index and predict a prediction interval compatible with literature data while avoiding the need for animal sacrifice (approximately 50 mice in the study by Sugihara *et al.*). Such findings had been previously suggested [10,13], but this is, to our knowledge, the first study to demonstrate them using a large panel of bacterial clinical isolates and a nonlinear mixed-effects pharmacodynamic model accounting for inter-strain variability.

Animal experiments are typically conducted on a limited number of strains, under the assumption that normalization of PK/PD indices by the MIC adequately accounts for potential inter-strain variability [1]. The *in vitro* approach proposed here enables PK/PD index determination across a large number of strains, thereby allowing direct quantification of inter-strain variability through nonlinear mixed-effects pharmacodynamic models. In our study, significant inter-strain variability was identified across several pharmacodynamic parameters, but this variability was observed across the whole range of strain susceptibility, *i.e.* the variability could not be explained by strain MIC, the PK/PD relationship was the same for low and high MIC strains. This unexplained-by-MIC inter-strain variability in the pharmacodynamic response to MEM translated into variability in the f%T>MIC values required to achieve a given bactericidal effect across strains. For instance, the 10th and 90th percentiles of f%T>MIC required to achieve a 2-log10 CFU reduction at 24 h were 27% and 71%, respectively. When bactericidal experiments are performed on only a few strains and analyzed in a pooled manner, the resulting PK/PD target tends to reflect the mean value. From a therapeutic perspective, however, it may be more appropriate to consider higher percentile targets—such as the 90th percentile (71%)—rather than the mean or median (44%), in order to better account for inter-strain variability. Thus, this in vitro framework may help mitigate biases associated with *in vivo* experiments conducted on a limited number of bacterial strains.

The quality of the *in vitro* pharmacodynamic model is critical for *in vivo* bacterial count simulations, as it defines the relationship between MEM concentrations and antibacterial effect. To our knowledge, this is the first study to use a nonlinear mixed-effects model to characterize inter-strain pharmacodynamic variability from in vitro bactericidal experiments. Bactericidal experiments were performed on a large collection of *Pseudomonas aeruginosa* clinical isolates (n = 66), covering a broad MIC range (≤0.5–32 mg/L) that included both susceptible and resistant isolates (EUCAST breakpoint = 8 mg/L). The structural model, combining an initial non-replicative state, an adaptive resistance mechanism, and MEM degradation, adequately captured both the initial delay in bacterial growth observed for some strains and subsequent bacterial regrowth. Random-effect parameters successfully described the observed inter-strain variability. The use of a sufficiently broad range of MEM concentrations was essential to appropriately characterize the concentration–effect relationship, particularly to identify whether an effect plateau (Emax) was reached. The existence of such a plateau indicates that, beyond a certain concentration threshold, further increases in drug exposure do not result in additional antibacterial activity, a hallmark of time-dependent antibiotics. Notably, simulated *in vivo* concentrations largely exceeded *in vitro* EC50 values, indicating that this plateau was frequently achieved under simulated conditions. This was necessary to confirm the time-dependent pharmacodynamic profile of MEM. We also identified a significant negative correlation between baseline EC50 (in the absence of adaptation) and the rate of adaptive resistance. This suggests that strains with greater initial susceptibility (lower EC50_ini_) tend to exhibit faster adaptive responses over time. It should be noted that other factors may influence PK/PD index targets, including immune status and bacterial inoculum size [1]. The impact of inoculum can be assessed *in vitro* through bactericidal experiments and incorporated into pharmacodynamic models [25]. By contrast, accounting for immune status remains more challenging. It is generally accepted that evaluating antibacterial activity under conditions lacking an immune response—such as *in vitro* bactericidal experiments or neutropenic mouse models—represents a worst-case scenario, as antibacterial efficacy would be expected to improve in the presence of host immune defenses.

*In vivo* simulations of bacterial counts also depend on the underlying PK model. The choice of PK model is therefore critical, as pharmacokinetics has been shown to influence the identification of PK/PD indices [10,15]. As discussed above, it is important to simulate sufficiently high concentrations to reach the effect plateau. However, simulations should remain within clinically relevant exposure ranges, as demonstrating time-dependent activity at supra-therapeutic concentrations would have limited translational value. In this study, we selected a murine PK model, adjusted for MEM protein binding in mice, to enable comparison with PK/PD results obtained in murine models. Nevertheless, the use of human PK models and clinically relevant dosing regimens may be more appropriate when aiming to determine PK/PD indices for clinical application [15]. In this context, population pharmacokinetic models are particularly useful, as they allow incorporation of inter-individual variability and ensure coverage of exposure ranges representative of those observed in clinical practice. It should also be noted that additional factors may influence PK/PD index targets, including the site of infection [1]. Drug exposure at the infection site can be accounted for using site-specific PK models when available [26].

In conclusion, we demonstrated that *in vitro* bactericidal experiments combined with nonlinear mixed-effects pharmacodynamic modelling not only allow identification of the PK/PD index associated with MEM efficacy, but also enable quantification of inter-strain variability. Replacing the traditional determination of PK/PD indices based on animal experiments with this *in vitro* approach warrants further investigation.

## Acknowledgments

We would like to thank Agnès Audurier for her excellent technical assistance.

## Author contributions

N.G.; V.A.-C. and J.M.B. designed the project. T.C. conducted experiments. O.K, N.G., and V.A.-C. conducted data analysis. N.G.; V.A.-C. and J.M.B. supported manuscript writing. All authors reviewed the manuscript, performed final editing, and approved the final version.

