## Supplemental text and figures for "Replacing *In Vivo* Experiments for PK/PD Target Determination Through *In Vitro* Time-Kill Experiments and PK/PD Modelling Incorporating Inter-strain Variability: Application to Meropenem Against *Pseudomonas aeruginosa*"

**Supplementary data**

List of supplements

Figure S1. Distribution of MICs for the 66 Pseudomonas aeruginosa clinical isolates used for *in vitro* Time-Kill experiments.

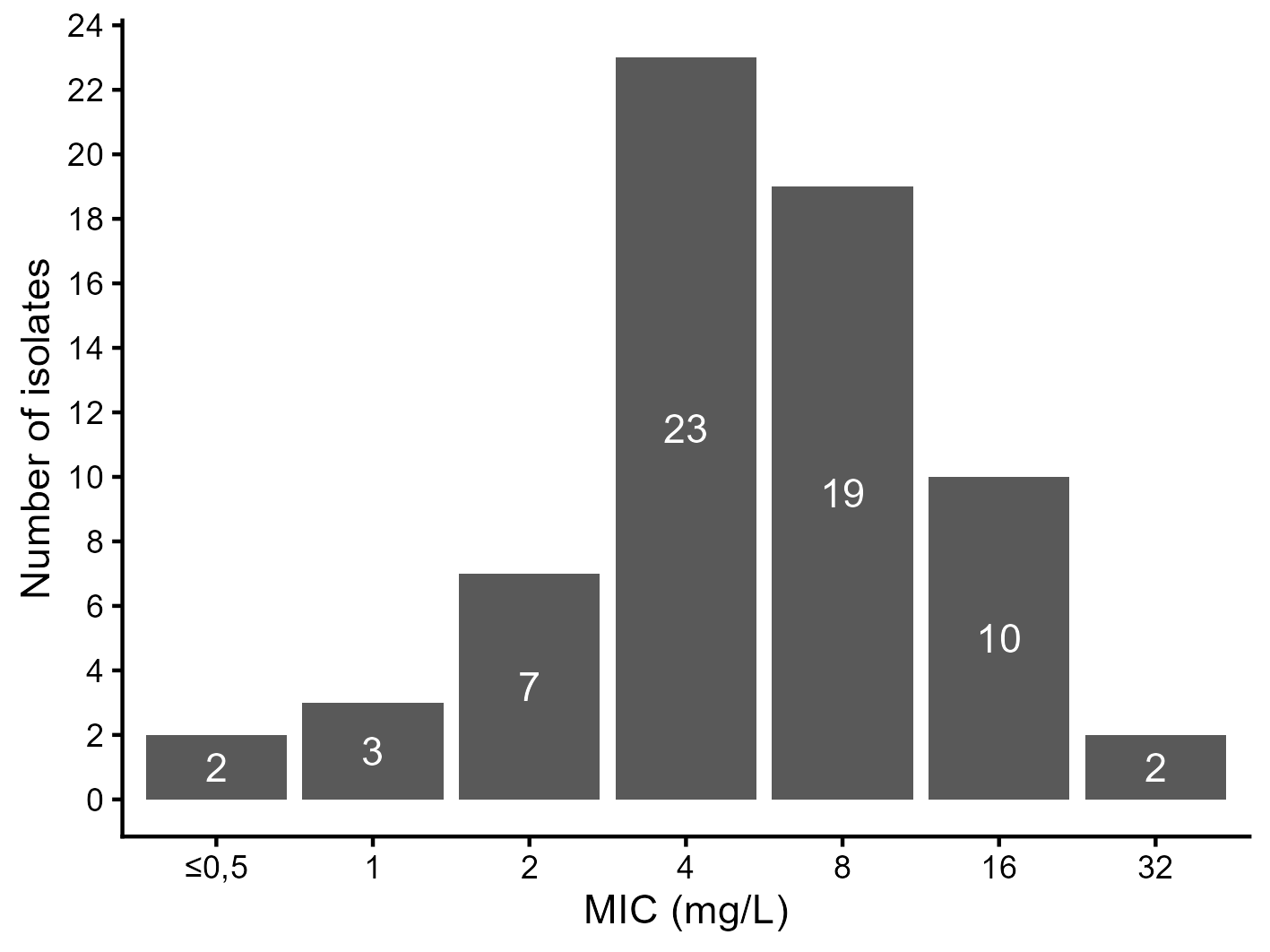

Figure S2. Individual plots for *in vitro* pharmacodynamic modelling. Points represent observed values, lines represent individual predictions, and values below the limit of quantification (log10 CFU = 3, red dashed line) are shown as bars.

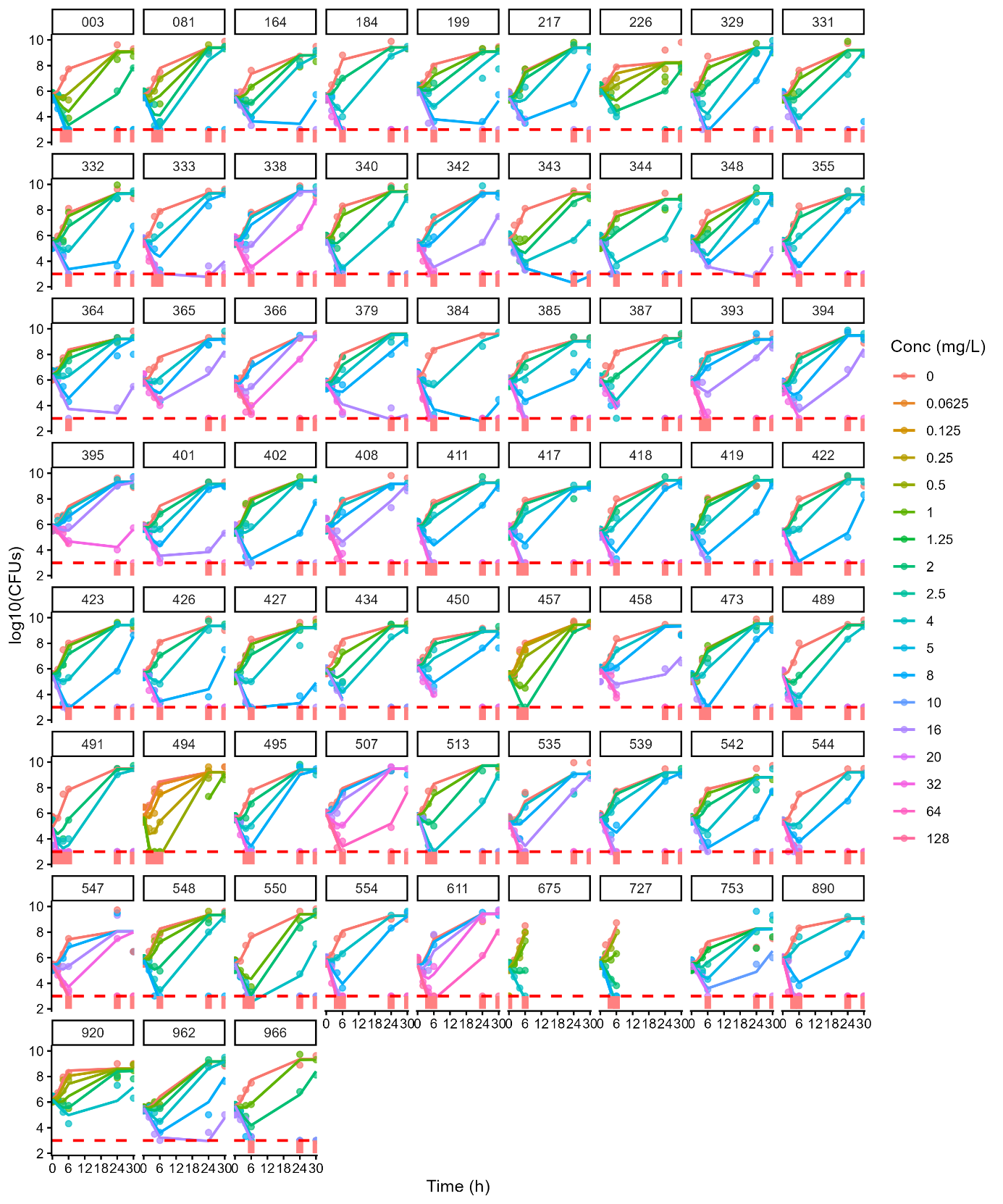

Figure S2. Normalized prediction distribution errors (NPDE) versus time and typical predictions. Pink triangles indicate NPDE for observations below the limit of quantification (based on model simulations).

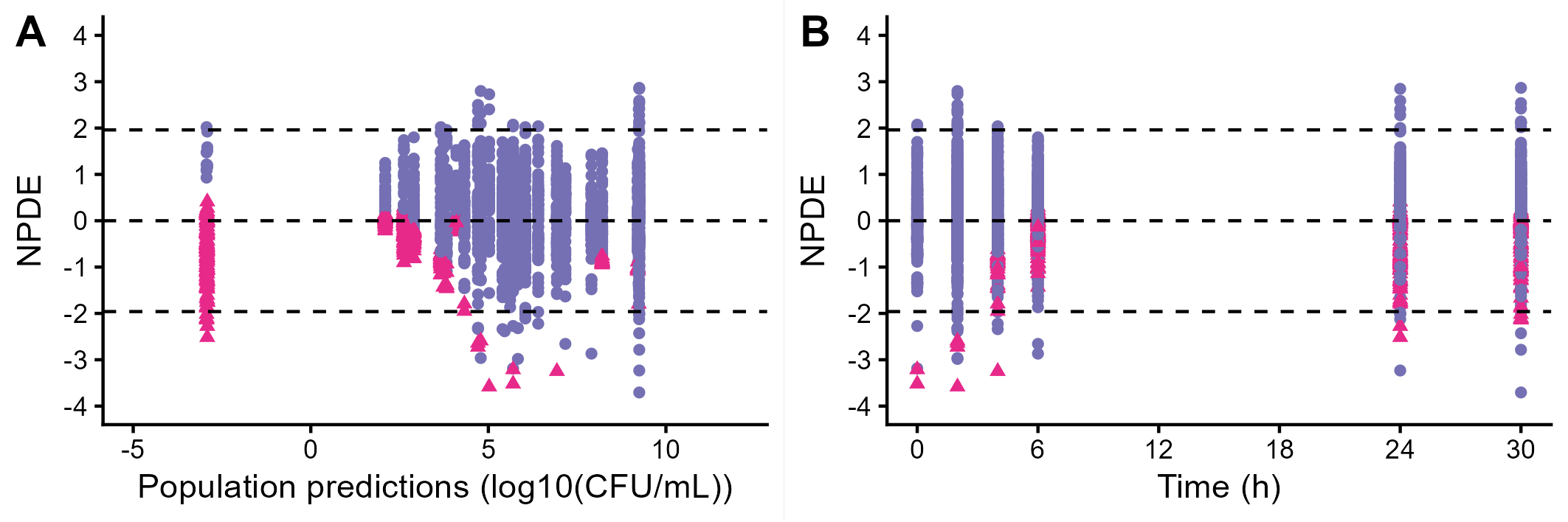

Figure S3. Observations versus typical population predictions.

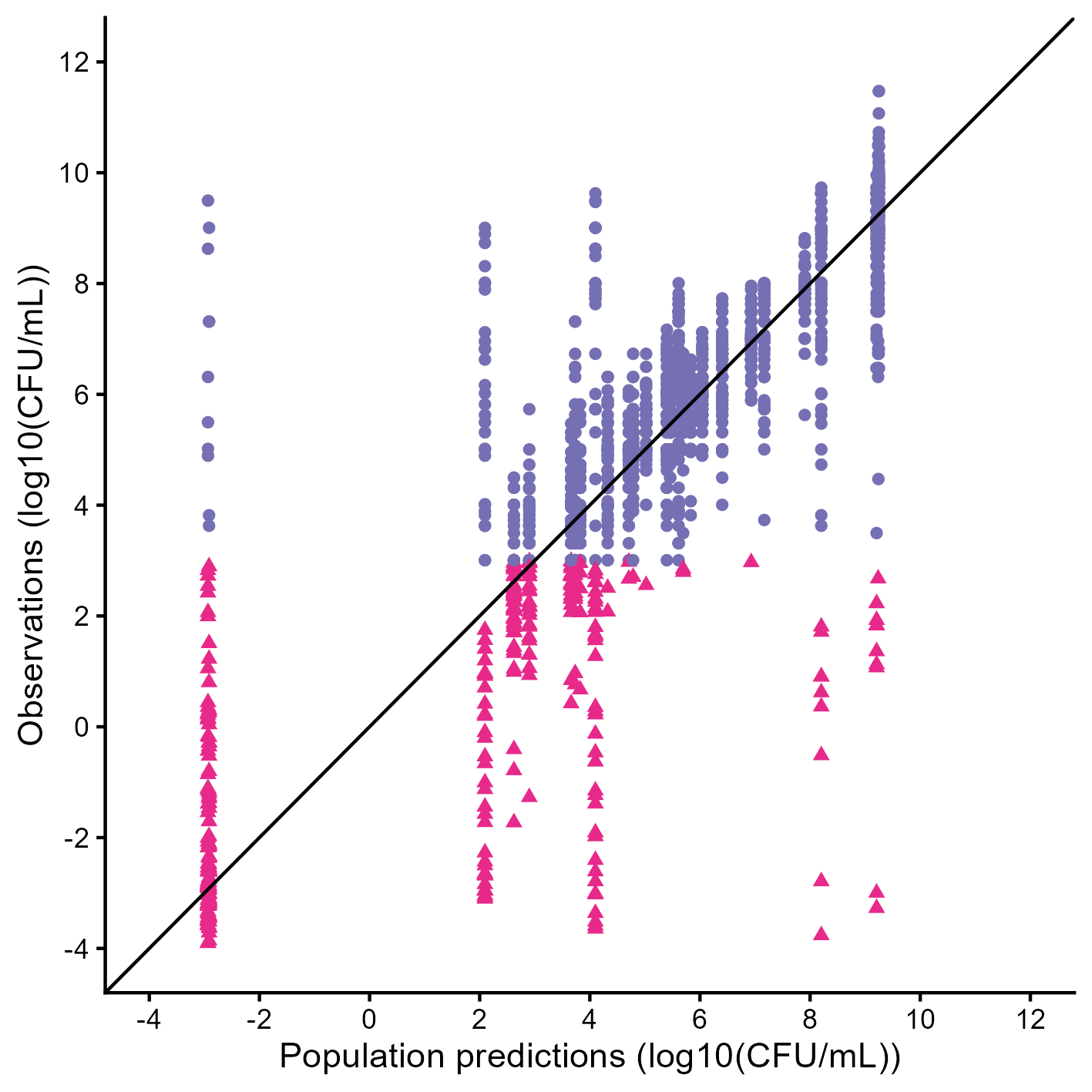

Figure S4. Observations versus individual predictions.

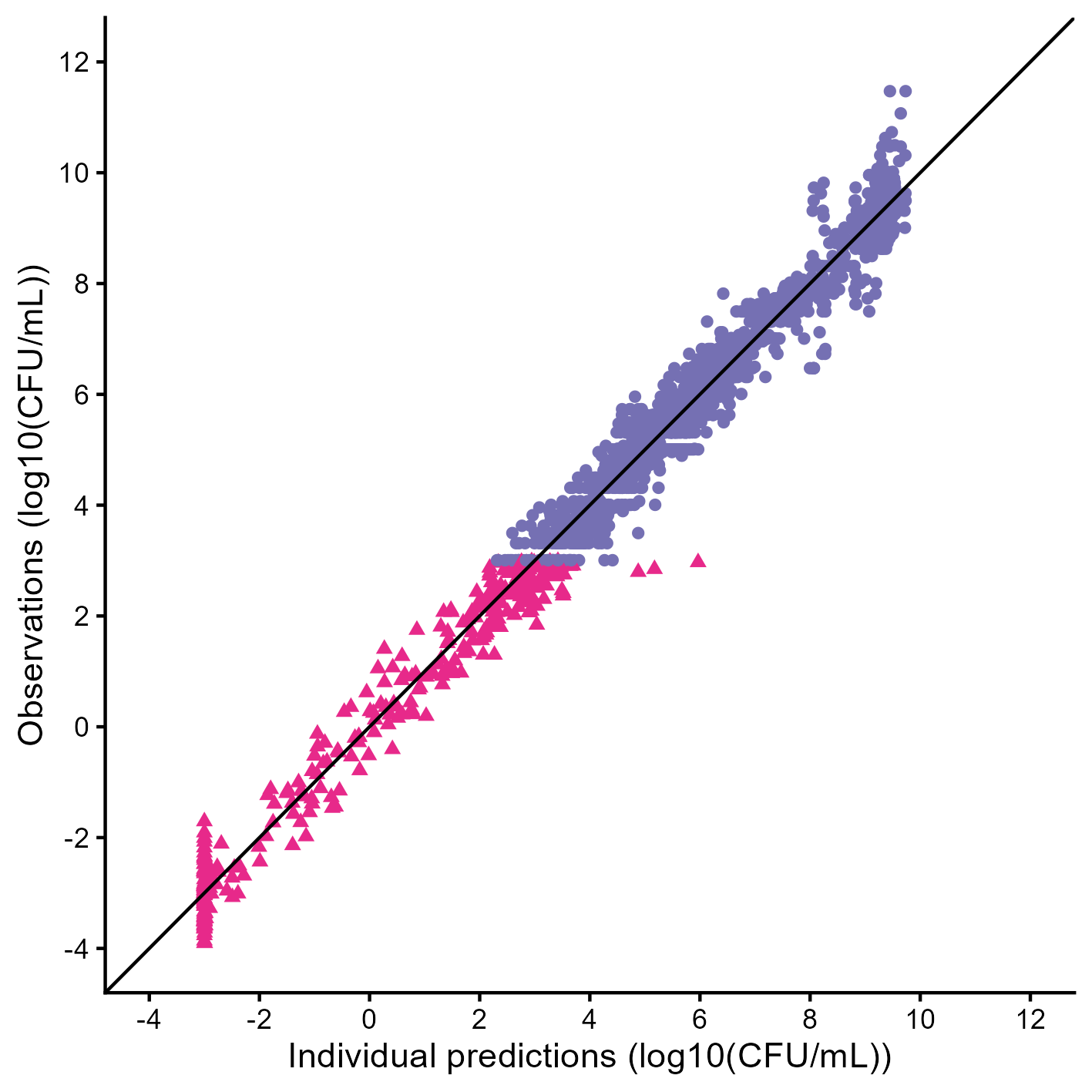

Figure S6. Median *in vivo* simulations of bacterial counts and plasma MEM concentrations following subcutaneous MEM dosing at 50, 100, 200, 400, and 800 mg/kg administered q24h, q12h, q6h, and q3h.

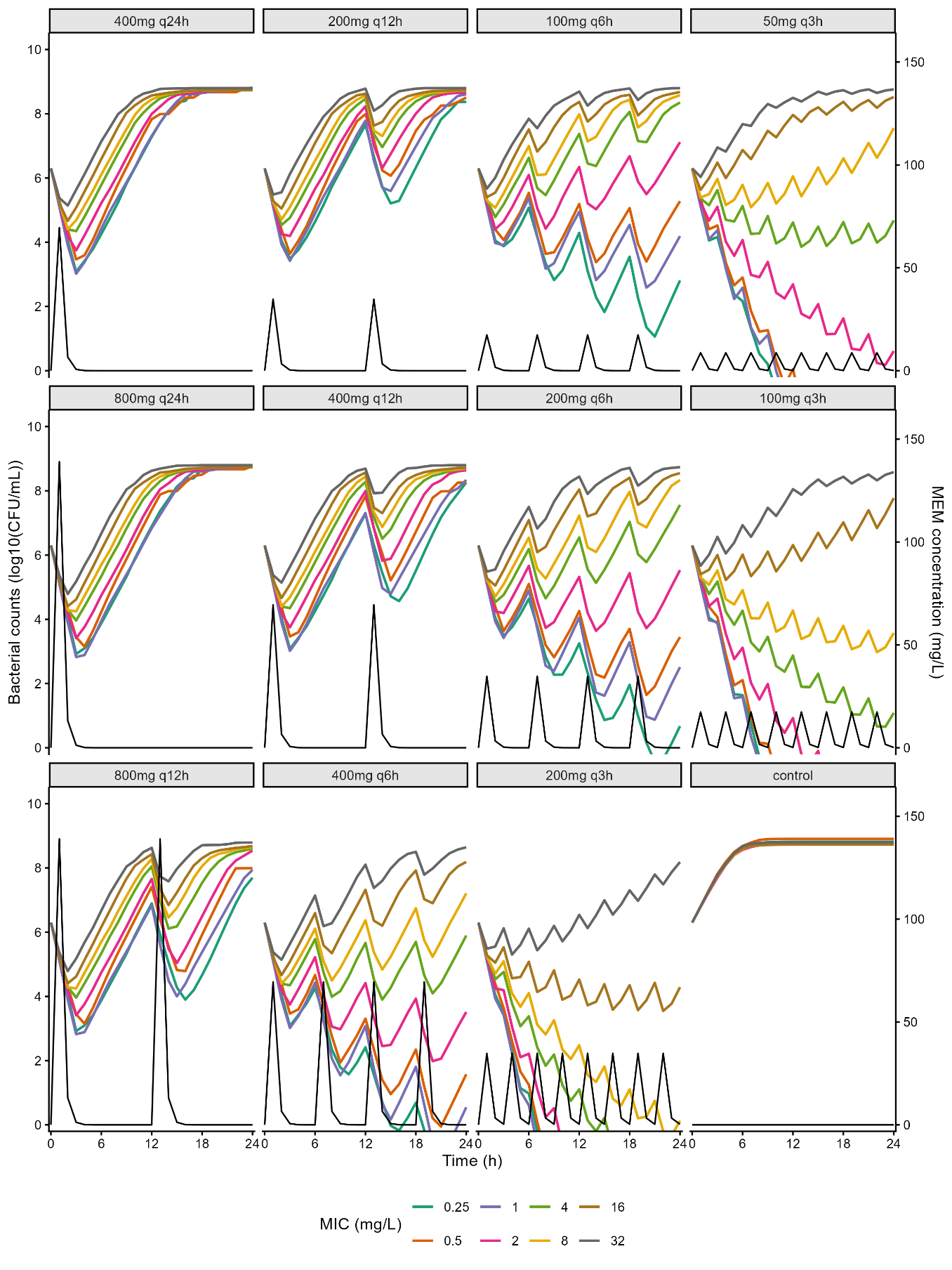

Table S1. Bootstrap results for the final *in vitro* pharmacodynamic model.

|  |  | Final model | | Bootstrap | |
| --- | --- | --- | --- | --- | --- |
| Parameter | Unit | Typical value | RSE (%)  Stochastic approx.. | Median | RSE(%) |
| f | Unitless | 0.28 | 4 | 0.27 | 13.6 |
| log10_Bmax | log_10_(CFU/mL) | 9.24 | 0.5 | 9.25 | 0.5 |
| k_net_ | h^-1^ | 1.15 | 2.5 | 1.16 | 6.6 |
| p | Unitless | 0.50 | 9.3 | 0.49 | 9.3 |
| k_RS_ | h^-1^ | 0.22 | 15.2 | 0.22 | 39.2 |
| Emax_S | h^-1^ | 3.58 | 4.9 | 3.80 | 10.3 |
| EC50ini | mg/L | 0.65 | 7.2 | 0.77 | 11.5 |
| γ | Unitless | 2.13 | 4.4 | 2.03 | 6.2 |
| Ada_EC50 | Unitless | 2.97 | 13.1 | 2.82 | 22.5 |
| γ_ada_ | Unitless | 3.51 | 8.8 | 5.95 | 25.3 |
| K_Ada_EC50_ | h^-1^ | 0.15 | 10.4 | 0.19 | 19.7 |
| omega_log10_Bmax | Log10(CFU/mL) | 0.34 | 10.8 | 0.34 | 19.6 |
| omega_knet | ln(h^-1^) | 0.14 | 13.8 | 0.13 | 38.7 |
| omega_p | Unitless | 0.67 | 10.1 | 0.68 | 12.2 |
| omega_k_RS_ | ln(h^-1^) | 0.80 | 15.7 | 0.87 | 26.2 |
| omega_Emax | ln(h^-1^) | 0.30 | 13.1 | 0.25 | 20.2 |
| omega_EC50ini | ln(mg/L) | 0.52 | 11 | 0.54 | 13.6 |
| omega_γ | Unitless | 0.53 | 14.3 | 0.50 | 20.6 |
| omega_Ada_EC50 | Unitless | 0.84 | 12.1 | 0.81 | 17.8 |
| omega_ γ_ada_ | Unitless | 1.35 | 21.9 | 1.80 | 38.2 |
| omega_Ada_EC50 | Unitless | 0.56 | 14.1 | 0.53 | 24.6 |
| corr_EC50_ini_ K_Ada_EC50_ | Unitless | -0.51 | 25.9 | -0.63 | 28.1 |
| a | log_10_(CFU/mL) | 0.44 | 1.8 | 0.44 | 4.3 |

Text S1. Pubmed queries to find *in vivo* meropenem PK/PD studies

**Query 1**

Date of last search: 2025-06-24
Query: murine thigh infection model AND meropenem AND pseudomonas aeruginosa

**Query 2**

Date of last search: 2025-06-24
Query: meropenem AND pseudomonas aeruginosa AND PKPD indices

**Query 3**

Date of last search: 2025-06-24
Query: meropenem AND pseudomonas aeruginosa AND mouse neutropenic infection model

**Query 4**

Date of last search: 2025-06-24
Query: meropenem AND pseudomonas aeruginosa AND dose fractionation study

Data and codes are available at the following address : <https://doi.org/10.57745/7UWQXI>

Data S1. Final TKC dataset

Data S2. PKPD targets collected from literature

Code S1. Monolix model for TKC analysis

Code S2. Simulx model for *in vivo* PKPD simulations

Code S3. Monolix code for nonlinear mixed effects regression of *in vivo* effect *versus* fT%>MIC

Code S4. Monolix code for nonlinear mixed effects regression of *in vivo* effect *versus* fCmax/MIC

Code S5. Monolix code for nonlinear mixed effects regression of *in vivo* effect *versus* fAUC/MIC
